# Photobiological Hydrogen Production at Scale: Integrating Bioprocess Optimization and Techno-Economic Modeling

**DOI:** 10.1101/2025.04.27.650233

**Authors:** Tamar Elman, Shabtai Isaac, Iftach Yacoby

## Abstract

Microalgal hydrogen production driven by solar energy offers significant promise as a sustainable energy alternative, yet remains economically challenging due to issues of scalability from laboratory to industrial applications. Here, we demonstrate scalable hydrogen production at semi-industrial volumes using the *Chlamydomonas reinhardtii pgr5* mutant, employing an optimized cultivation protocol and photobioreactor design. This approach achieves a fivefold increase in hydrogen yield. Notably, the post-production biomass maintains a high-quality protein and nutrients profile, emphasizing microalgae as a “green coin” with energy security on one side and food security on the other. Techno-economic analysis suggests, achievable hydrogen production costs could reach $2.70/kg H_2_ under projected improvements, and potentially decrease furthur to $1.48/kg with full optimization. By effectively bridging laboratory research and practical industrial implementation, our study establishes a dual-purpose algal hydrogen production platform aligned with circular economy principles, positioning microalgae prominently within sustainable energy and food frameworks.

## Main

Gaseous hydrogen (H_2_) is considered an ideal energy source due to its high energy density, dual role as both an energy carrier and a fuel, and its zero emissions at the point of use ^1^. However, since H_2_ is scarcely abundant on Earth, it must be extracted from natural compounds such as water. Current production methods – primarily based on fossils fuels are classified using a color code that reflects their carbon footprint: black H_2_ denotes the most polluting methods, while green H_2_, produced using renewable energy sources, aims for net-zero emissions ^2^. Green H_2_ is predominantly generated via electrolysis, a process in which water is split into oxygen (O₂) and H_2_ using electricity, with reported efficiencies reaching up to 80% ^3^. Nevertheless, producing 1 kilogram of H_2_ requires 42-70 kWh of energy—far exceeding its usable energy output of 33 kWh^4^. This unfavorable energy return, along with additional challenges inherent to green H_2_ production, poses substantial economic barriers to competing with fossil fuels ^5^. Electrolysis’s high energy demand is attributed to the strong chemical bonds between hydrogen and oxygen atoms in water molecules, which require substantial input to break. This bond is naturally cleaved only through photosynthesis - a biological process that harnesses solar energy to convert water and carbon dioxide into organic molecules, releasing O₂ as a byproduct ^6^. A subset of photosynthetic organisms, including specific bacteria and microalgae, can produce H_2_ as a byproduct at the end of the photosynthetic electron transport chain, catalyzed by the enzyme Hydrogenase (HydA) ^7^. HydA functions as a “safety valve”, releasing excess electrons to prevent system overload ^8,9^. Consequently, its activity is short-lived and disrupted by two factors: the diversion of electrons toward carbon fixation, and the accumulation of O₂, which irreversibly inactivates the enzyme ^10^. Over recent decades, advances in strain engineering and cultivation methods have extended the H_2_ production phase from minutes to days and even weeks ^11–22^. However, two critical challenges remain in the field. First, scalability: most academic studies operate at small volumes, and their results are frequently extrapolated without adequate verification. Indeed, previous studies that proactively tested scalability at larger volumes (50–100 liters) encountered significant non-linearities and reduced yields compared to small-scale experiments ^23–25^. Second, like many renewable technologies, algal H_2_ production remains costly to operate and maintain, particularly when compared to the low market price of H_2_, which is expected to remain competitive with fossil-derived sources. Together, these challenges hinder the transition from promising laboratory findings to practical and economically viable applications.

To address these challenges, we developed a systematic approach to scale up H_2_ production from microalgae, transitioning from laboratory-scale to semi-industrial volumes. Utilizing the *Chlamydomonas reinhardtii* (*C.r*) *pgr5* mutant strain -known for its ability to sustain continuous H_2_ production ^26–28^, we introduced iterative engineering refinements alongside adaptations to the cultivation and H_2_ production protocol. Real-time gas quantification enabled accurate assessment of H_2_ output, thereby improving both precision and reliability. Through optimization of photobioreactor design and key process parameters, we overcame expected scalability limits, achieving a fivefold increase in H_2_ yields over our previous benchmarks ^22^.

Furthermore, our production protocol ^21^, in contrast to the widely used sulfur-deprivation protocol, maintains algal biomass integrity post-H_2_ production ^29^. This distinction significantly enhances the economic potential of the system, allowing the residual biomass to serve as a valuable secondary product that helps offset operational costs. Compositional analysis confirmed the nutritional quality of the post-H_2_ production biomass, supporting its applicability across a range of market sectors. Coupled with additional features like carbon fixation and dark cultivation, which provide both economic and environmental benefits, we conducted a techno-economic analysis to evaluate an integrated model for concurrent biomass and H_2_ production. The analysis was based on an existing data stream from a commercial algae production company and incorporated results from the present study. We examined the integration of algal cultivation and H_2_ production systems under varying growth conditions (e.g., culture density) and H_2_ production rates. The projected costs were further evaluated across different types of cultivation facilities, with the goal of identifying operational parameters that could support economic feasibility. Taken together, this work demonstrates the successful volumetric scaling of algal H_2_ production, and establishes a foundational economic framework for its practical implementation in existing agricultural and industrial contexts.

### Direct scale-up of H_2_ production results in non-linear and reduced performance

To scale up published results tenfold, we constructed an 11-liter tubular photobioreactor (PBR; 10 L culture, 10 cm diameter; Fig. 1a), using the same tools and methods previously applied in small - scale laboratory experiments ^21^. All PBRs were illuminated with “cool white” LEDs (Fig. 1c) and aerated during biomass accumulation. Mixing was maintained by magnetic stirring or peristaltic pumping to ensure homogeneity throughout anaerobic H_2_ production. H_2_ output was measured directly from the reactor headspace using volumetric systems. Cultures were initiated by a 1:10 dilution of mid-log phase cells (15–25 μg Chl mL^−1^) and grown under continuous light and air flow. Once the cultures reached 30–35 μg Chl mL^−1^ (∼1 g dry weight L^−1^), the H_2_ production protocol was applied ^21^. A single gas chromatography (GC) measurement from the 11-liter PBR’s headspace verified that the small-scale H_2_ production phenotype was retained at larger volume (Fig. 1b). Following this validation, all subsequent measurements were performed by gas volume only, using various volumetric devices. Over one week, the 10-liter culture yielded 956 mL H_2_— only 13.6% of the expected 7 liters based on linear extrapolation from 1-liter cultures ^22^. This discrepancy underscores the challenges of scale-up, likely caused by heterogeneity in larger volumes. Variations in light distribution, gas exchange, and mixing reduced productivity, emphasizing the need for direct volumetric H_2_ measurement at scale.

**Figure 1.**
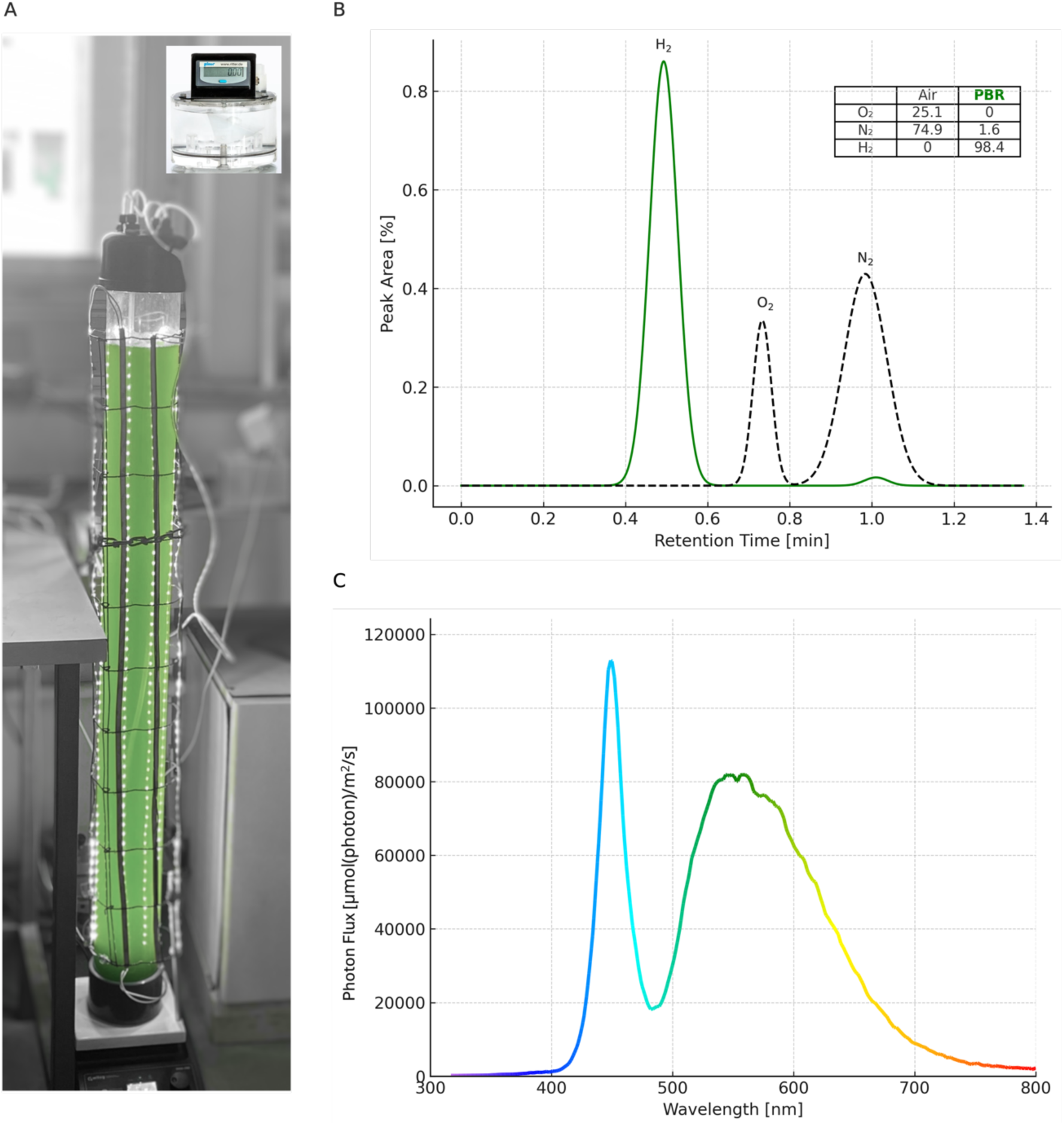
11 L Tubular PBR Setup for H_2_ Production. (**a**) PBR (10 cm diameter) containing 10 L of *pgr5* culture (30 μg Chl mL^−1^) under ∼400 μE m^−2^ s^−1^ light. The cap includes an air inlet (for growth) and a gas outlet connected to a Ritter MilliGascounter (inset). (**b**) GC analysis showing H_2_ as the main gas in the reactor headspace (solid line), compared to ambient air (dashed line). Percent composition is shown in the side table. (**c**) Spectral distribution of 6500K LEDs used for both cultivation and hydrogen production.

### Fine Engineering of Photobioreactor Design Leads to 5-fold Increase in H_2_ Production Efficiency

To address these challenges, we evaluated how reactor geometry influences H_2_ production efficiency and identified the optimal design for further process development. Two Plexiglas (poly(methyl methacrylate)) PBR prototypes (4 L each) were constructed: one retained the familiar tubular structure (9 cm diameter, Fig. 3b; green rectangles), while the other explored a panel-type design with a 4 cm light path (Fig. 2b). The panel PBR’s short light path enhances penetration, and its high surface-area-to-volume ratio and modular design make it well-suited for scale-up. Unlike the tubular PBR, which used magnetic stirring during H_2_ production, the panel model required external circulation via a peristaltic pump, continuously moving culture in and out of the panel’s narrow corners (Fig. 2b, side view).

**Figure 2:**
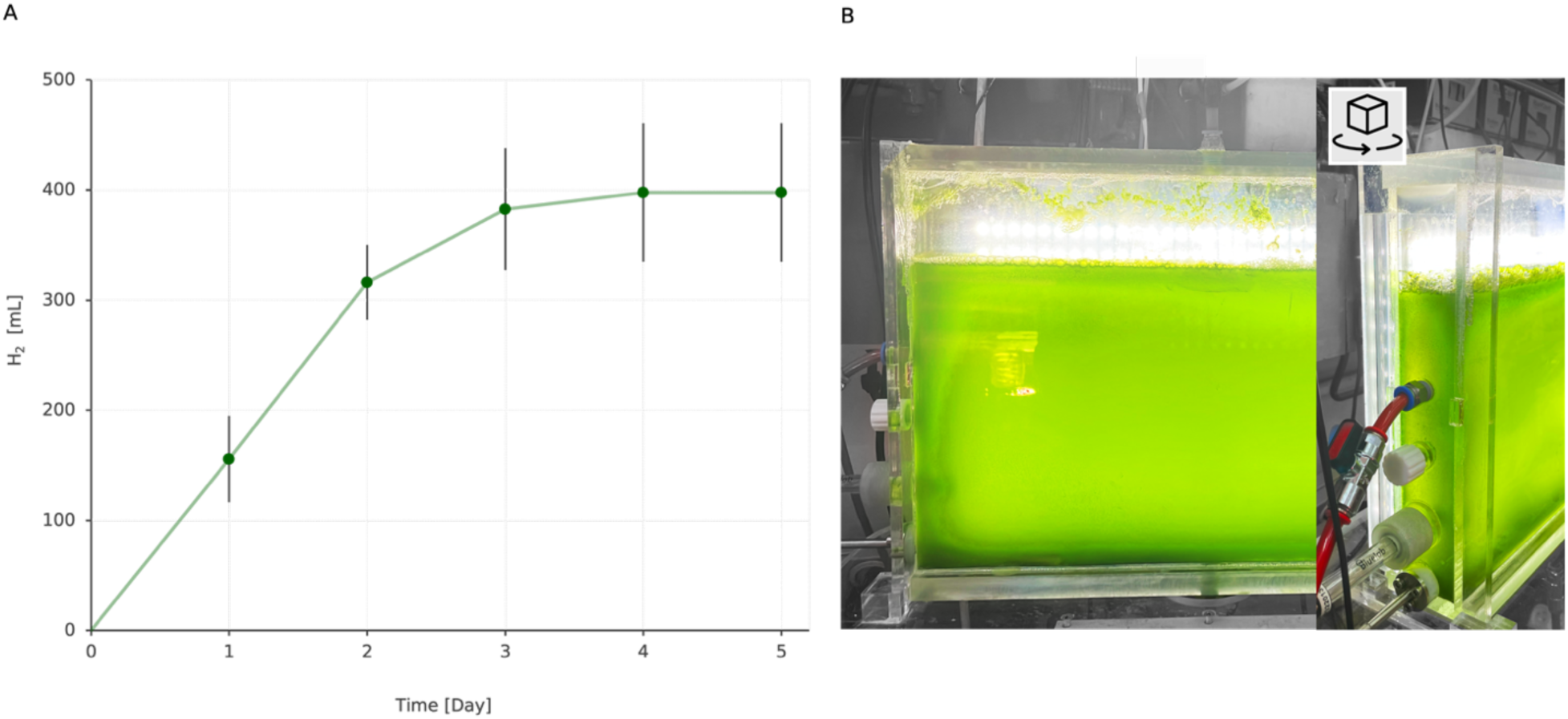
H_2_ production in a flat-panel photobioreactor. (**a**) H_2_ accumulation [mL] over five days in a flat-panel PBR containing *pgr5* cultures (3 L) at 30 μg Chl mL^−1^, continuously illuminated at ∼400 μmol photons m^−2^ s^−1^. Mean values ± standard error are shown (n = 3). (**b**) Experimental setup with LED illumination, a gas outlet (top section) connected to collection systems (BlueVis Count), and real-time data logging (BlueVis software). The right panel details reactor inlets, including a peristaltic pump, an air inlet for culture mixing, and a pH monitoring probe.

**Figure 3.**
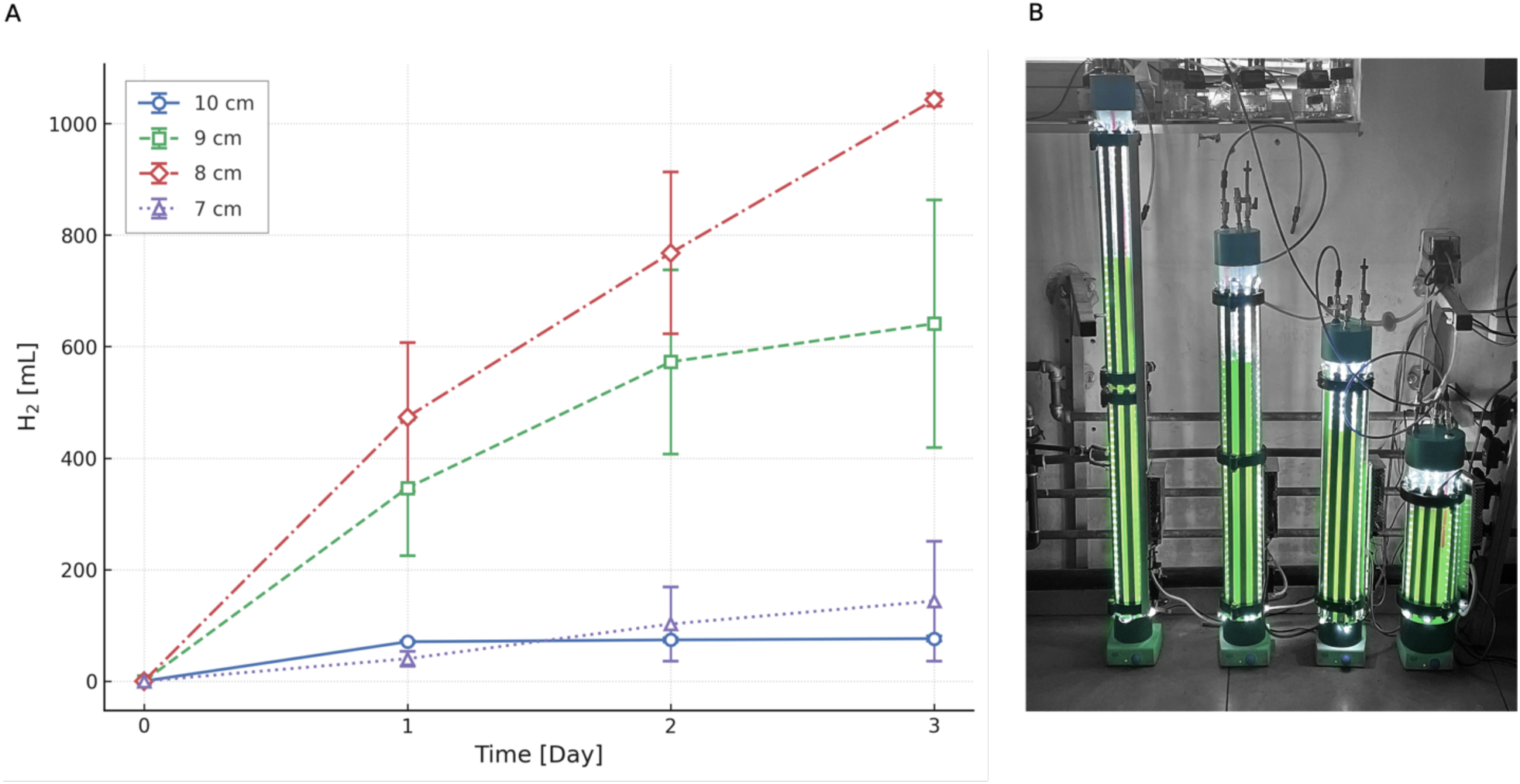
H_2_ production in 3 L tubular PBRs of varying diameters. (**a**) H_2_ accumulation [mL] over three days in tubular PBRs with diameters of 10 cm (blue), 9 cm (green), 8 cm (red), and 7 cm (purple). *pgr5* cultures (3 L) at 30 μg Chl mL^−1^ were continuously illuminated at ∼400 μmol photons m^−2^ s^−1^. Mean values ± standard error are shown (n = 3). (**b**) Experimental setup with LED illumination, gas collection via Ritter MilliGascounter, and real-time data logging (BlueVis software). Reactor diameters increase from left (7 cm) to right (10 cm).

In a single day, the 9 cm tubular PBR generated an amount of H_2_ as equivalent to that produced by the panel PBR over a five-day period (Fig. 3a, green rectangles; Fig. 2a), demonstrating a clear efficiency discrepancy. While both systems had comparable light paths, the panel PBR’s external circulation likely reduced mixing efficiency and introduced geometry-related limitations affecting gas behavior and productivity. Although panel PBRs offer scalability advantages, improving mixing is critical to unlocking their full potential for H_2_ production.

To further investigate these parameters, we expanded the tubular PBR model to include four reactors with diameters of 7, 8, 9, and 10 cm (Fig. 3b). By maintaining a constant culture volume of 3 L across all diameters, we assessed how changes in light penetration, mixing, and liquid column pressure—each influenced by reactor geometry—affected H_2_ production and its transition from the soluble to gaseous phase. The results were striking: the 8 cm PBR (red diamonds) produced 1 liter more H_2_ than the 10 cm PBR (blue circles) over four days. While the 10 cm reactor exhibited reduced light permeability, the large difference in yield suggests that light intensity is a key driver of H_2_ production efficiency. However, the 7 cm PBR (purple triangles), despite its higher light transmission, produced similar amounts of H_2_ as the 10 cm unit, indicating the influence of additional factors. This unexpected trend points to two possible explanations: (1) excessive light intensity in smaller reactors may impair the photosynthetic machinery, reducing H_2_ output; or (2) taller liquid columns may increase hydrostatic pressure, affecting gas exchange and limiting H_2_ release. This added pressure, along with suboptimal mixing, may delay H_2_ degassing. It is also likely that these parameters act synergistically, collectively shaping overall yields.

Having identified the optimal reactor type and dimensions, we replicated the design and incorporated a specialized cap with a pH meter inlet and a feed tube, enabling controlled titration to maintain constant acidity (Fig. 4b). Designed specifically for the H_2_ production phase, the cap allowed external probe access while preventing H_2_ leakage. The feed-on-demand system employed is a simplified, low-tech adaptation of fed-batch methods commonly used in agriculture and industrial fermentation. These systems support precise biological control at scale by dynamically adjusting conditions and nutrient input based on culture needs.

**Figure 4.**
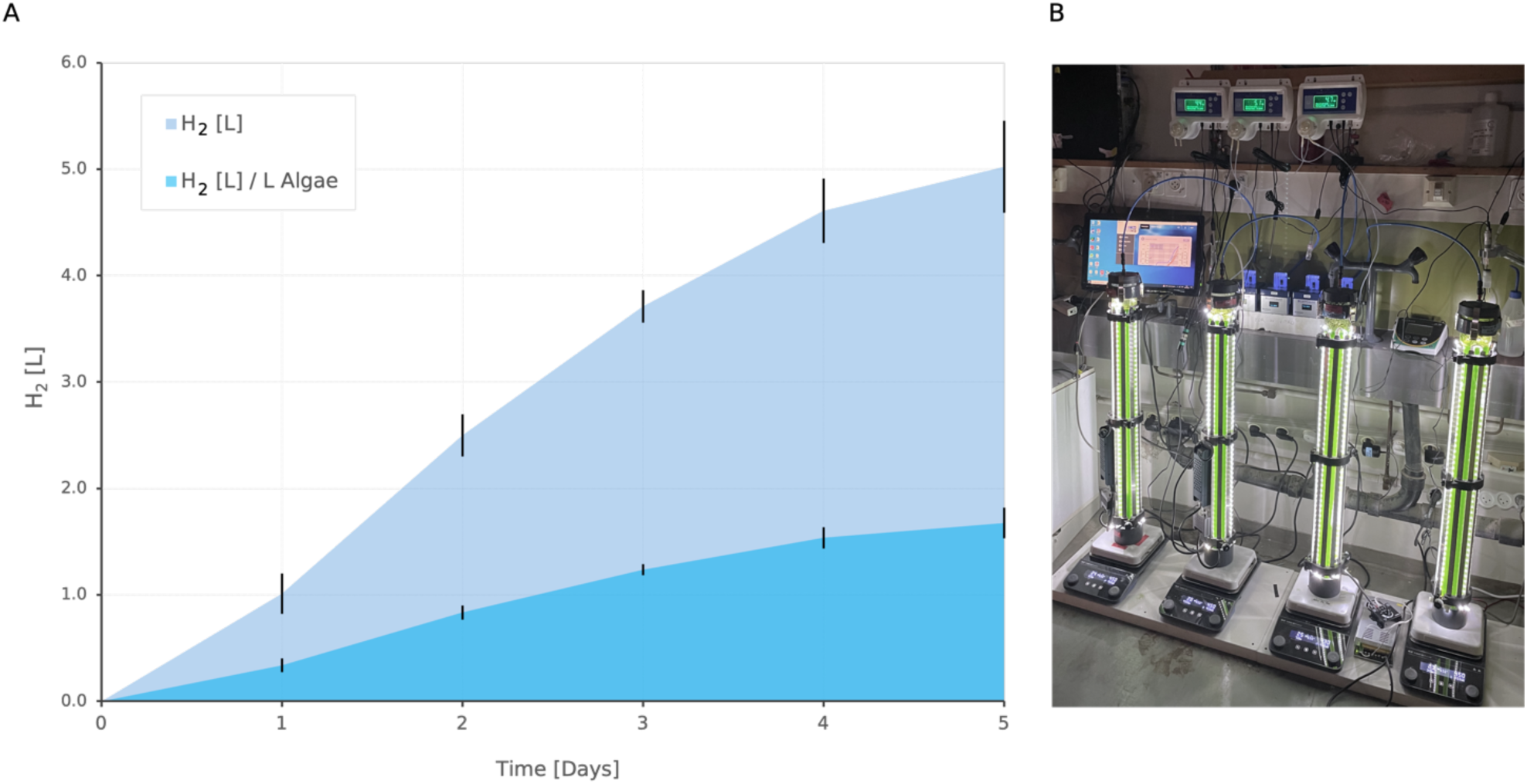
H_2_ production in 8 cm tubular photobioreactors. (**a**) Total H_2_ production (L) and normalized H_2_ productivity (L H_2_ per L algae) over five days in tubular PBRs. *pgr5* cultures (3 L) at 30 μg Chl mL^−1^ were continuously illuminated at ∼400 μmol photons m^−2^ s^−1^. Mean values ± standard error are shown (n = 5). (**b**) Experimental setup showing the PBRs with LED illumination, gas collection systems (BlueVis Count), and real-time data logging (BlueVis). The reactor cap includes a gas outlet and an inlet for continuous pH monitoring with automatic acetate feedback addition (Bluelab pH Controller).

In our setup, acetate—an external carbon source for microalgae—served dual roles: supporting mitochondrial respiration to remove oxygen and acting as a reductant for the photosynthetic system, both contributing to sustained H_2_ production ^30^. Over five days, the 3-liter culture accumulated an average of 5 liters of H_2_, equivalent to 1.5 liters per liter of algal culture (Fig. 4a). By comparison, using the previously reported protocol ^22^, a 1-liter culture produced 700 mL of H_2_ over 12 days. This reflects a substantial improvement, with the new setup generating 300 mL/day versus 58 mL/day—more than a fivefold increase.

Previous attempts to produce H_2_ using *C. reinhardtii* in mid-scale photobioreactors as well as our preliminary attepmt (Fig. 1) showed the challanges of scaling up small-scale results ^24^. Owning to that, we carried out matrice of adaptations, allowing our system to achive higher yields. This achievement stems from a combination of optimized protocols, reactor modifications, physiological enhancements, and the selection of an appropriate algal strain—together enabling a more effective transition from laboratory scale (mL) to semi-industrial volumes (liters). The system also provides strong evidence for the feasibility of scaling up to larger, more advanced cultivation platforms.

### Dual Utility Focus: Unlocking the Value of Post-H_2_ Algal Biomass

Traditional approaches to algal H_2_ production rely on sulfur deprivation to induce hypoxia and activate the oxygen-sensitive HydA enzyme ^31^. While effective at triggering H_2_ evolution, this method is inefficient, requires frequent medium replacement, and causes severe physiological stress and cell death—limiting both its scalability and the downstream utility of the resulting biomass. In this study, we sought to overcome these limitations by developing an integrated model that supports H_2_ production under ambient conditions, while also enabling the recovery of high-quality algal biomass suitable for diverse industrial applications. To evaluate the viability of our ambient protocol, we performed a comprehensive nutritional analysis of *pgr5* biomass post-H_2_ production. Cell survival and biomass retention were first assessed, yielding an average of 0.87 g dry weight (dw) L^−1^, with cultures retaining a vital appearance (Fig. 5a–b). Given its relevance for food and feed applications, dried biomass was analyzed at Mérieux NutriSciences for macronutrient content (Fig. 5c). The results revealed a high protein content (47%), along with fats (10%) and carbohydrates (28%), amounting to 392 kcal per 100 g. Amino acid profiling (Fig. 5d, S1) confirmed that *pgr5* exceeds the Dietary Reference Intake (DRI) for key amino acids ^32^ highlighting its potential as a high-quality protein source. Fat analysis (Fig. 5e, Table S1) showed mono-unsaturated (3.4 g/100 g dw) and polyunsaturated (1.3 g/100 g dw) fatty acids, with a favorable omega-3 to omega-6 ratio of 2:1 ^33^. Notably, palmitic acid (36.66%) and stearic acid (10.15%) comprise a major share of the lipid profile, suggesting potential for biofuel applications due to their stability and energy density (Table S1). Carotenoid profiling revealed substantial levels of lutein (930 mg/kg dw) and beta-carotene (661 mg/kg dw), both valued for their antioxidant properties and vision-supporting roles ^34^ (Fig. 5e). Together, these findings emphasize the versatility of *pgr5* biomass, supporting its viability for nutritional and industrial uses beyond H_2_ production. In parallel, we conducted a detailed characterization of the biomass’s micronutrient profile, including essential minerals, vitamins, and heavy metals. The results, provided in Supplementary Table S2, further underscore the nutritional quality and safety of the biomass for food-grade and industrial applications.

**Figure 5.**
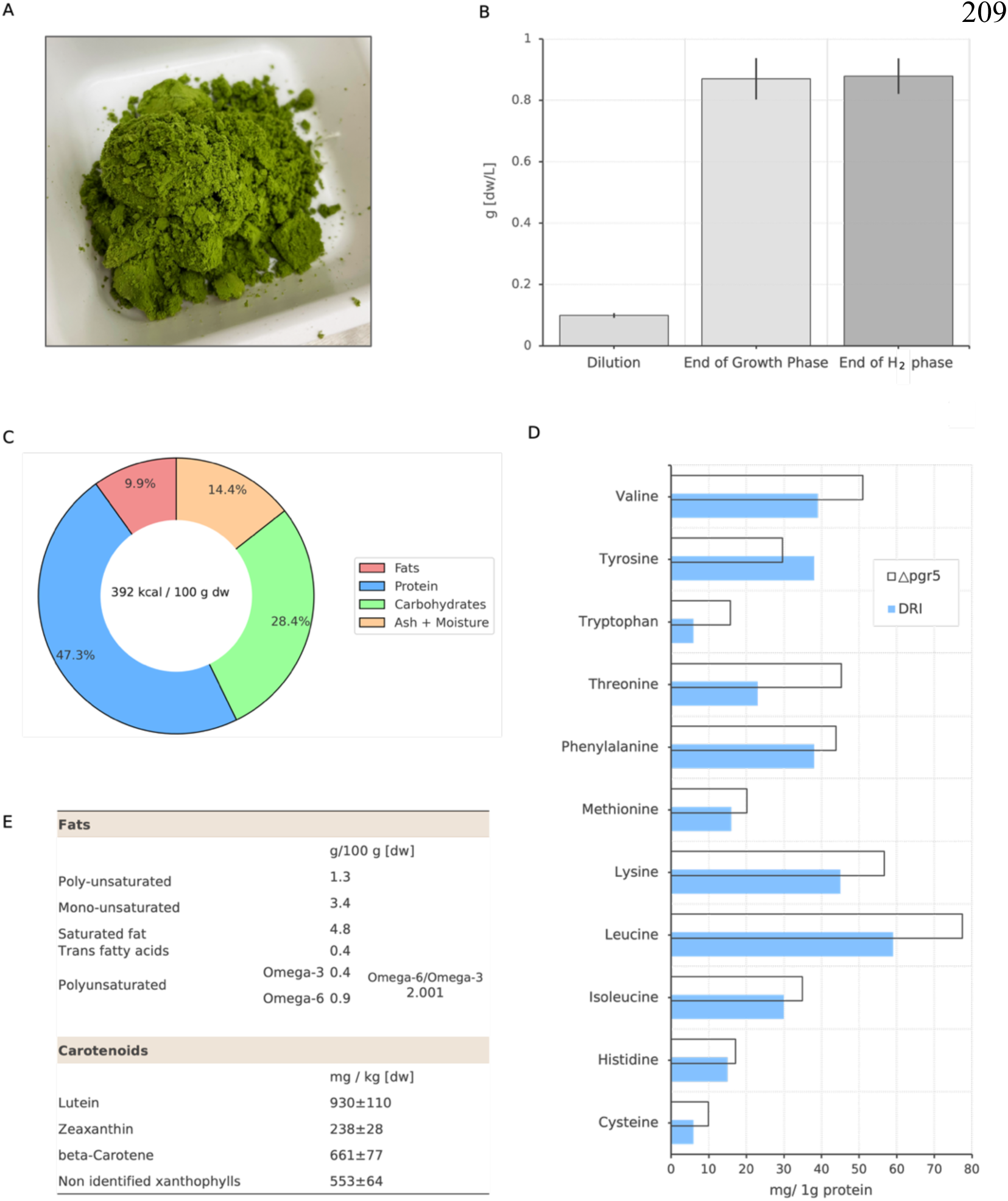
Composition and nutritional profile of *pgr5* biomass after H_2_ production. (**a**) Dried biomass at the end of the H_2_ production phase. (**b**) Biomass concentration (g/L dry weight) measured at three stages: initial growth, post-growth, and post-H_2_ production. Mean values ± standard error are shown (n = 9). (**c**) Macronutrient composition, including fat, protein, carbohydrate percentages, and caloric value. (**d**) Essential amino acid profile (mg per 1 g protein) in comparison to dietary reference intake (DRI). (**e**) Detailed breakdown of fat composition (g per 100 g dry weight) and carotenoid content (mg per kg dry weight).

Taken together, the measured nutritional and biochemical components align with previously reported values for the *pgr5* background strain *C. reinhardtii* ^35^, while also revealing the accumulation of valuable compounds. This consistency confirms that our ambient H_2_ production protocol preserves biomass integrity and quality, reinforcing the dual utility of *pgr5* as both a sustainable H_2_ production platform and a nutrient-rich resource for food, feed, and nutraceutical markets.

### Balancing Carbon Capture and Energy Production

To align with the Intergovernmental Panel on Climate Change (IPCC) greenhouse gas reduction targets ^36^, both emerging technologies and established companies are pursuing strategies to optimize carbon offsetting. These efforts aim to reduce pollution penalties—anticipated to increase under future regulations—while maximizing the economic value of carbon credits through trading or sale.

In this context, we evaluated modifications to our system that integrate environmental and economic benefits. To assess the carbon assimilation potential of *pgr5* during biomass growth, we monitored CO₂ concentrations under aerobic conditions using Membrane Inlet Mass Spectrometry (MIMS). Cultures were exposed to alternating light and dark cycles with continuous aeration (Fig. 6). The average carbon fixation rate during illuminated phases was –12 µmol CO₂/hr^-1^/mg Chl^-1^. This corresponds to 0.39 g CO₂ fixed per gram dry weight of algae per day—comparable to reported rates for various microalgae species ^37^. These results confirm that under aerobic conditions, *pgr5* effectively fixes carbon, reinforcing its potential contribution to carbon credit generation.

**Figure 6.**
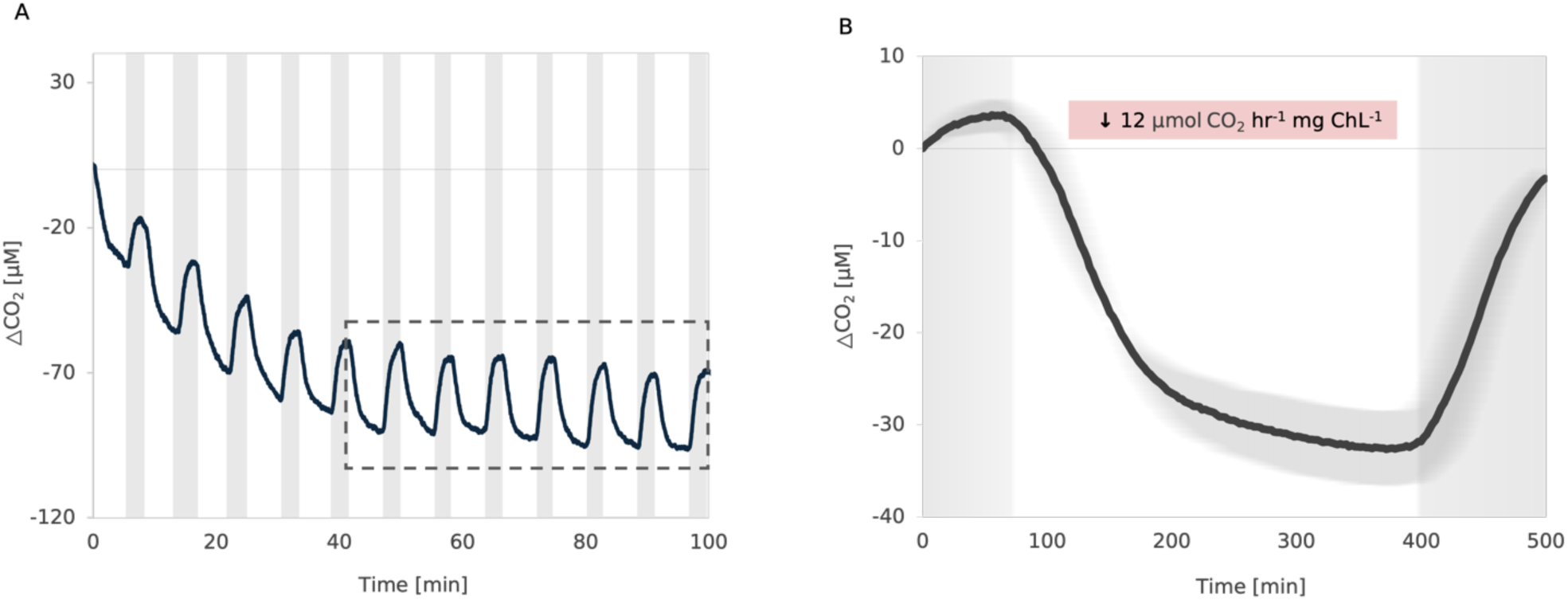
Aerobic Carbon Fixation in *pgr5*. (**a**) Dissolved CO₂ dynamics in *pgr5* cultures monitored by membrane inlet mass spectrometry (MIMS) at 45 µg Chl mL^−1^ under red actinic light (1500 µmol photons m^−2^ s^−1^). Cultures (5 mL) were continuously aerated to maintain aerobic conditions and subjected to 12 alternating light–dark cycles (dark phases shaded). The dashed box indicates the seven cycles averaged in panel (b). (**b**) Averaged CO₂ response across the final seven cycles. Data are presented as mean ± standard error (n = 3).

We further explored the integration of dark growth phases as a strategy to reduce cultivation costs and enhance carbon offset potential. Dark cultivation modifies cellular metabolism, potentially influencing downstream products such as H_2_ upon re-illumination. As our system aims to combine energy production with cost-effective biomass generation, we sought to determine whether algae grown in the dark could still support efficient H_2_ production.

In our experiments, *pgr5* was grown in 1-liter cultures under dark conditions until the target biomass concentration was reached. The cultures were then transitioned to light, and H_2_ production was monitored (Fig. 7). Over seven days, the culture produced approximately 530 mL of H_2_— within the average range reported for grown-illuminated *pgr5* cultures ^22^.

**Figure 7.**
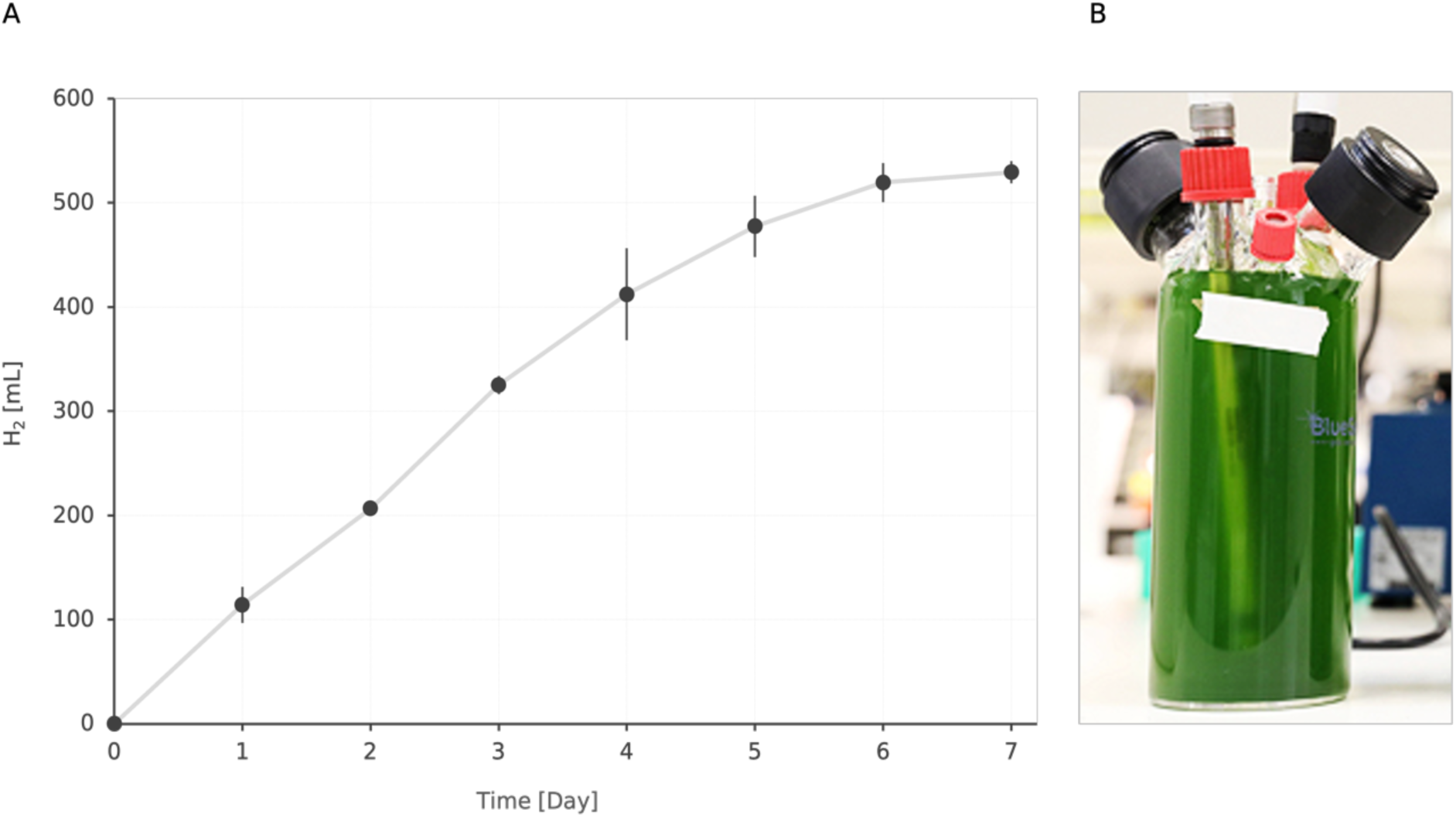
Algal H_2_ production after dark mixotrophic growth. (**a**) Cumulative H_2_ production (mL) over 8 days in 1-L *pgr5* cultures initiated at 8 μg Chl mL^−1^ and exposed to continuous illumination (∼400 μmol photons m^−2^ s^−1^, n = 2). (**b**) Experimental setup using a 1-L BlueSens photobioreactor for biomass growth and H_2_ monitoring. Mid-log phase cells (∼5 μg Chl mL^−1^) were diluted into 950 mL of fresh medium and grown under dark conditions until reaching 10–15 µg Chl mL^−1^, at which point the H_2_ production protocol was applied.

This experiment demonstrates the feasibility of applying the H_2_ production protocol to dark-grown biomass. Future work will aim to optimize dark cultivation parameters under controlled conditions to minimize performance gaps relative to light-grown cultures and facilitate the integration of dark phases into scalable production systems.

### Techno-economic Analysis

To perform the techno-economic analysis, we selected “Bar Algae”, a profitable microalgae company, specializing in biomass sales —as a representative case study. We assessed the feasibility and economic impact of retrofitting their facility to support integrated algal biomass and H_2_ production. This adaptation involved capital expenditures (CAPEX) for H_2_ compression and storage infrastructure, as well as operational expenses (OPEX) including energy and labor.

The model was based on a 1.2-hectare facility with a total culture volume of 720 m³. As shown in Figure 4, the observed average H_2_ production rate was 0.35 L/day (19.88 µmol H_2_/mg Chl/hr), with a peak of 0.5 L/day (28.40 µmol H_2_/mg Chl/hr) on day two. Two production scenarios were considered: one using the measured average rate under current lab conditions (1 g/L), and one projecting future high-performance systems sustaining peak production. To achieve a daily yield of 1 kg H_2_, 31 kg of algal biomass would be required at the average rate, or 22 kg at the peak rate (Table S3).

The corresponding levelized cost of hydrogen (LCOH)—which accounts for both upfront CAPEX and ongoing OPEX—was estimated at $12.94–17.62 per kg H_2_ (Fig. 8a). LCOH reflects the net present value of total production costs distributed over the facility’s operational lifetime and total H_2_ output.

**Figure 8.**
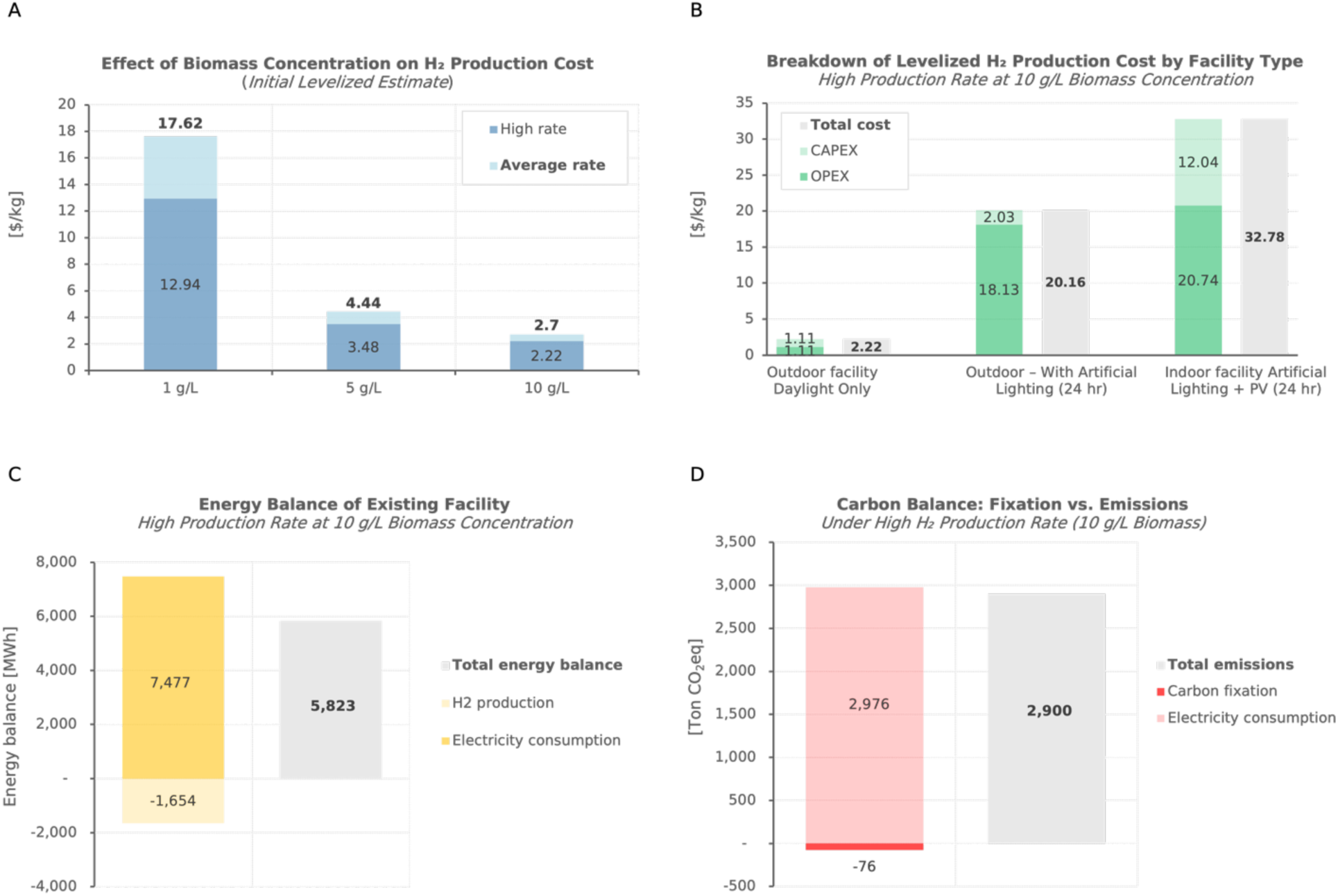
Techno-economic analysis of H_2_ production in a converted microalgae cultivation facility. **(a)** Levelized H_2_ production costs ($/kg H_2_) under average and peak production rates at biomass concentrations of 1, 5, and 10 g/L. **(b)** Production costs ($/kg H_2_) associated with three facility configurations at 10 g/L biomass and high H_2_ production rate: (i) existing outdoor facility with compression and storage systems; (ii) same with added LED lighting for continuous illumination; (iii) indoor facility with artificial lighting and photovoltaic (PV) integration. **(c)** Energy balance (MWh/year) of the facility under high production rate and 10 g/L biomass. **(d)** Carbon emissions (t CO₂eq/year) under high production rate and 10 g/L biomass, including contributions from electricity consumption and carbon fixation.

We further analyzed the impact of increasing biomass concentration on production economics. While the initial system was modeled at 1 g/L biomass concentration, we evaluated scenarios with 5 and 10 g/L—reflecting achievable targets for controlled large-scale cultivation systems^38^ . At 5 g/L, the estimated LCOH dropped to $3.48–4.44 per kg H_2_, and at 10 g/L it further decreased to $2.22–2.70 per kg H_2_, for the high and average production rates respectively (Fig. 8a).

Building on these findings, we modeled three potential adaptation scenarios for high-rate H_2_ production at 10 g/L biomass concentration (Fig. 8b):

1. Minimal adaptation of the existing outdoor facility, incorporating H_2_ compression and storage.
2. Addition of LED lighting to enable continuous illumination (daylight plus artificial light at night).
3. Conversion to a fully indoor cultivation system with artificial lighting and rooftop photovoltaic (PV) panels to reduce thermal regulation energy costs.

While scenarios 2 and 3 offer 24-hour operation and greater environmental control, they also significantly increased production costs to $20–32 per kg H_2_ (Fig. 8b). These increases were driven by both the capital investment in infrastructure and, more critically, by the high electricity demands of artificial lighting—outweighing the energy savings from PV supplementation and reduced thermal losses.

Considering the demonstrated improvement in H_2_ yields from photobioreactor optimization and the reported *in vivo* maximum rate of ∼120 µmol H_2_/mg Chl/hr ^39^, we explored a conservative future scenario with a threefold increase in H_2_ productivity. Under such optimized conditions, the projected LCOH could be reduced to $1.32–1.48 per kg H_2_ (Fig. S2).

Finally, we evaluated the potential contribution of H_2_ production to the facility’s overall energy and carbon footprint. Given hydrogen’s energy content of 33 kWh/kg, it could offset over 22% of the electricity consumed by the existing algae facility (Fig. 8c). However, direct carbon mitigation via algal carbon fixation remained modest compared to emissions from electricity use—especially when sourced from non-renewable energy (Fig. 8d).

## Conclusions

This study set out to answer two fundamental questions: (1) Can successful laboratory-scale results of algal H_2_ production be effectively scaled up? and (2) From a critical techno-economic perspective, is such an effort worthwhile?

The first question received a clear and encouraging answer. We achieved a fivefold scale-up of H_2_ production relative to our original laboratory setup, reaching days-long, stable H_2_ production at approximately one-quarter of the theoretical maximum photosynthetic rate (29 µmol H_2_ mg[chl]^−1^ h^−1^). This performance exceeded our expectations and demonstrates that photobiological H_2_ production is not inherently limited to small-scale experiments. The improved H_2_ production rates reported in this study were achieved solely through engineering adjustments to the photobioreactor design. This underscores the significant untapped potential of dedicated engineering development tailored to H_2_ production processes.

The answer to the second question is more nuanced. From a purely economic standpoint, H_2_ production alone becomes feasible only at high biomass concentrations—conditions that still require further demonstration. When placed within the broader context of a profitable, operational microalgae biomass facility, H_2_ production presents additional advantages. It can serve as an auxiliary product stream, adding value beyond the primary biomass output. Alternatively, it can function as an internal buffering system, capturing and storing energy already invested in biomass growth, and potentially reducing the overall energy footprint of the facility.

Overall, our findings suggest that while algal H_2_ is unlikely to serve as a primary energy source for heavy industry, it aligns well with the needs and constraints of biomass-oriented microalgae platforms or local, independent energy production facilities, as our analysis confirms that biomass quality is not compromised during the H_2_ production phase. In fact, the biomass is always worth more than the H_2_ it produces—yet this very relationship creates an opportunity: the biomass stream can effectively subsidize H_2_ production, enabling algal H_2_ to reach highly competitive pricing. This work lay the groundwork for implementing algal H_2_ production as a viable addition to existing microalgae industries, offering a flexible model for facilities seeking to diversify outputs and enhance energy efficiency.

## Methods

### Microalgal Strains and Cultivation Protocols

The working stock of *Chlamydomonas reinhardtii* mutant strain *pgr5* was cultivated in Tris– Acetate–Phosphate (TAP) medium under continuous agitation in Erlenmeyer flasks. The cultures were maintained at a constant temperature of 24°C with an irradiance of approximately 60 µmol m^−2^ s^−1^. The day before MIMS experiments, cultures were diluted into fresh TAP to ensure that they reached an early logarithmic growth phase (5-7 mg Chl mL^−1^) the following day. For experiments conducted in photobioreactors with a volume exceeding one liter, where sterilization of the growth medium by autoclaving was not feasible, a sterilization protocol based on tap water treatment with bleach and ascorbic acid was implemented as described in ^40^. Chlorophyll content was extracted and quantified following the protocol established by Jeffrey and Humphrey ^41^.

### MIMS Measurements

Membrane Inlet Mass Spectrometry (MIMS) analysis was conducted following the methodology outlined in ^42^. To assess CO_2_ levels, 5 mL of culture containing 45 µg Chl mL^−1^ in TAP medium, supplemented with 50 mM HEPES (4-(2-hydroxyethyl)-1-piperazineethanesulfonic acid), was transferred into a quartz cuvette and positioned within a chamber (Optical unit ED-101US/MD, Walz). To maintain aerobic conditions, the culture was aerated using a steady stream of air from a pump. After a 10-minute adaptation period in darkness to equilibrate CO_2_ levels, the samples were exposed to red actinic light at an intensity of 1,500 µmol m^−2^ s^−1^ for 5 minutes, using a Dual-Pulse Amplitude Modulated Fluorometer (DUAL-PAM-100; Heinz Walz Gmbh, Effeltrich, Germany), followed by a 2.5-minute dark phase. This sequence was repeated for 12 cycles and averaged for CO_2_ fixation trend. CO_2_ traces were normalized using calibration factors derived from standard curves, as detailed in Liran et al ^42^.

### H_2_ production assays

All H_2_ production experiments were conducted as outlined in ^21^. The dark 1-L experiment was carried out in a 1.15-L BlueSens bioreactor, while larger volume experiments utilized Perspex photobioreactors with varying geometries and diameters. Each photobioreactor was equipped with a cool white 6500K LED lighting system to provide an irradiance of 400 µmol m^−2^ s^−1^ in water. H_2_ outputs were measured using a Ritter MilliGascounter or BlueVcount gas counters (BlueSens), and with real-time data acquisition via Bluevis software (BlueSens) when indicated. If mentioned, pH monitoring was maintained using Bluelab pH Controller. During biomass accumulation, culture mixing was sustained through aeration using an air pump, while in the H_2_ production phase, mixing was facilitated by either a magnetic stirrer (VELP Scientifica or Witeg) or a peristaltic pump (BioRad) for the panel photobioreactor.

### Biomass Composition Analysis

Following the H_2_ production phase, the culture was centrifuged at 7,000 g for 10 minutes. The remaining biomass was concentrated into pellets in 50 mL tubes and immediately frozen at -80°C. For sample drying, the pellets were immersed in liquid nitrogen for 10 minutes and then transferred directly into a Labcono freezone Lyophilizer at a constant temperature of -80°C under a vacuum of minimum 0.12 bar for 24 hours. The dried biomass was subsequently used for analysis. The compositional analysis of microalgal biomass was outsourced to an external certified laboratory, Mérieux NutriSciences Corporation, at Milouda & Migal Laboratories, Acre, Israel. These analyses were conducted using industry-standard methods. Detailed methods and data supporting these analyses are available in the Source Data files.

### Technoeconomic analysis

The analysis of the levelized cost of H_2_ production was based on the assessed lifecycle CAPEX and OPEX of converting an existing facility. A yearly interest rate of 10% and a system life period of 25 years were considered for the analysis. The required systems were sized based on the calculations and assumptions detailed in Table S3, Source Data Table S3 and the following table:

### Equations and parameters used in techno-economic analysis

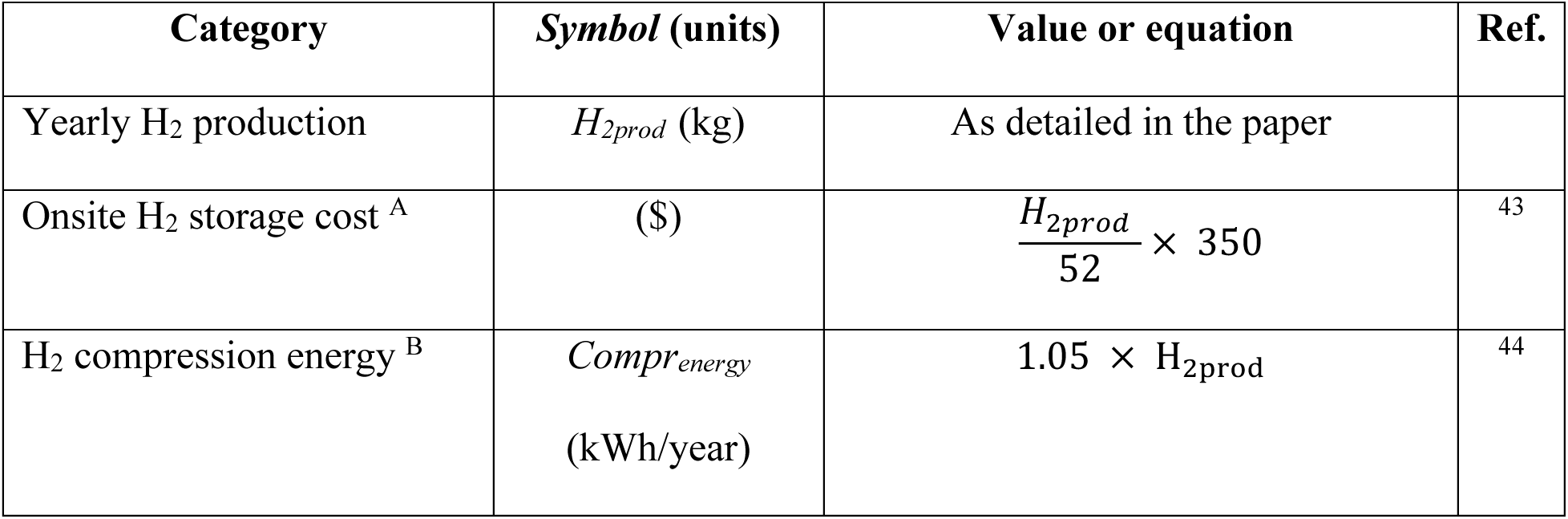
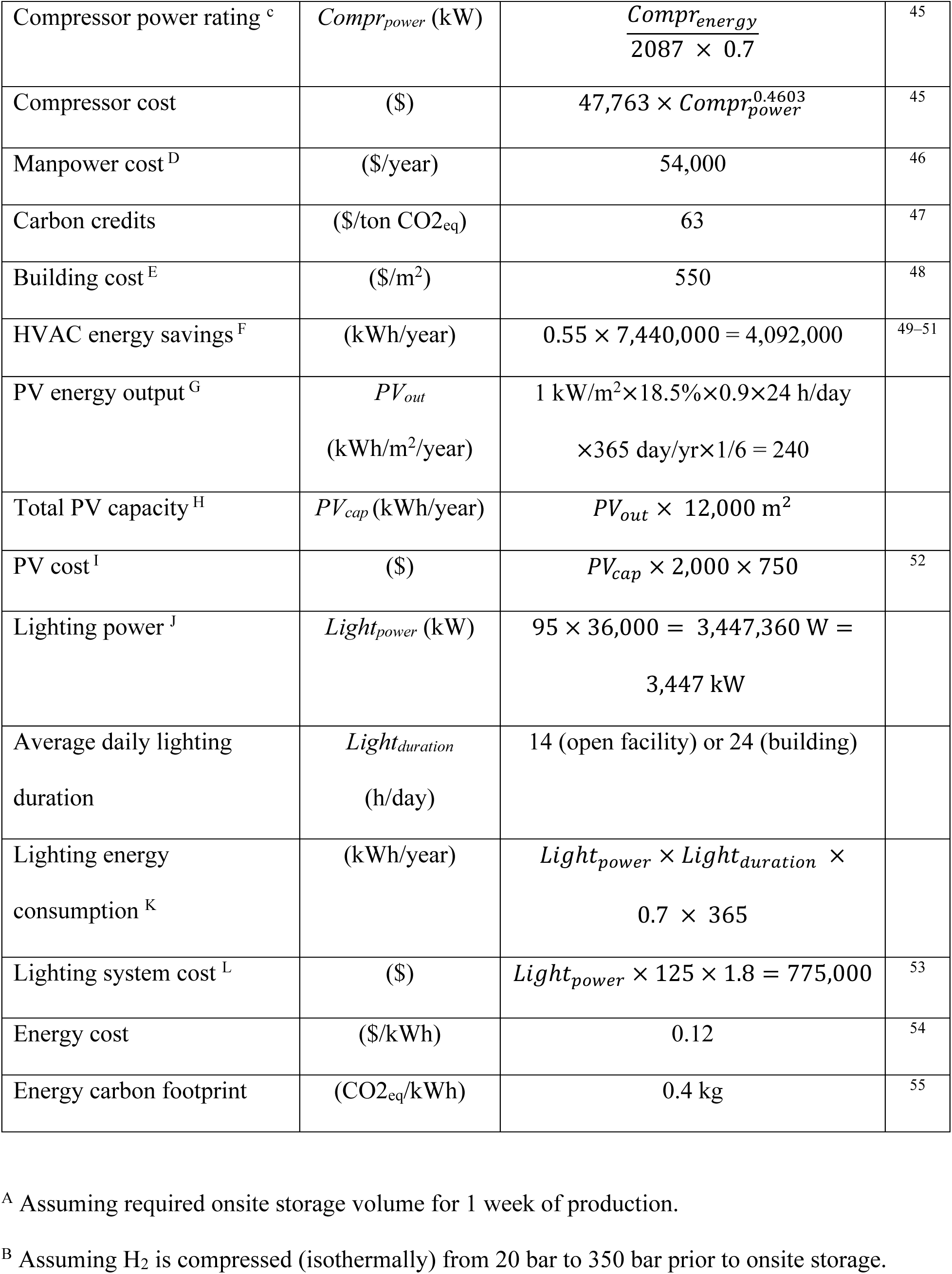
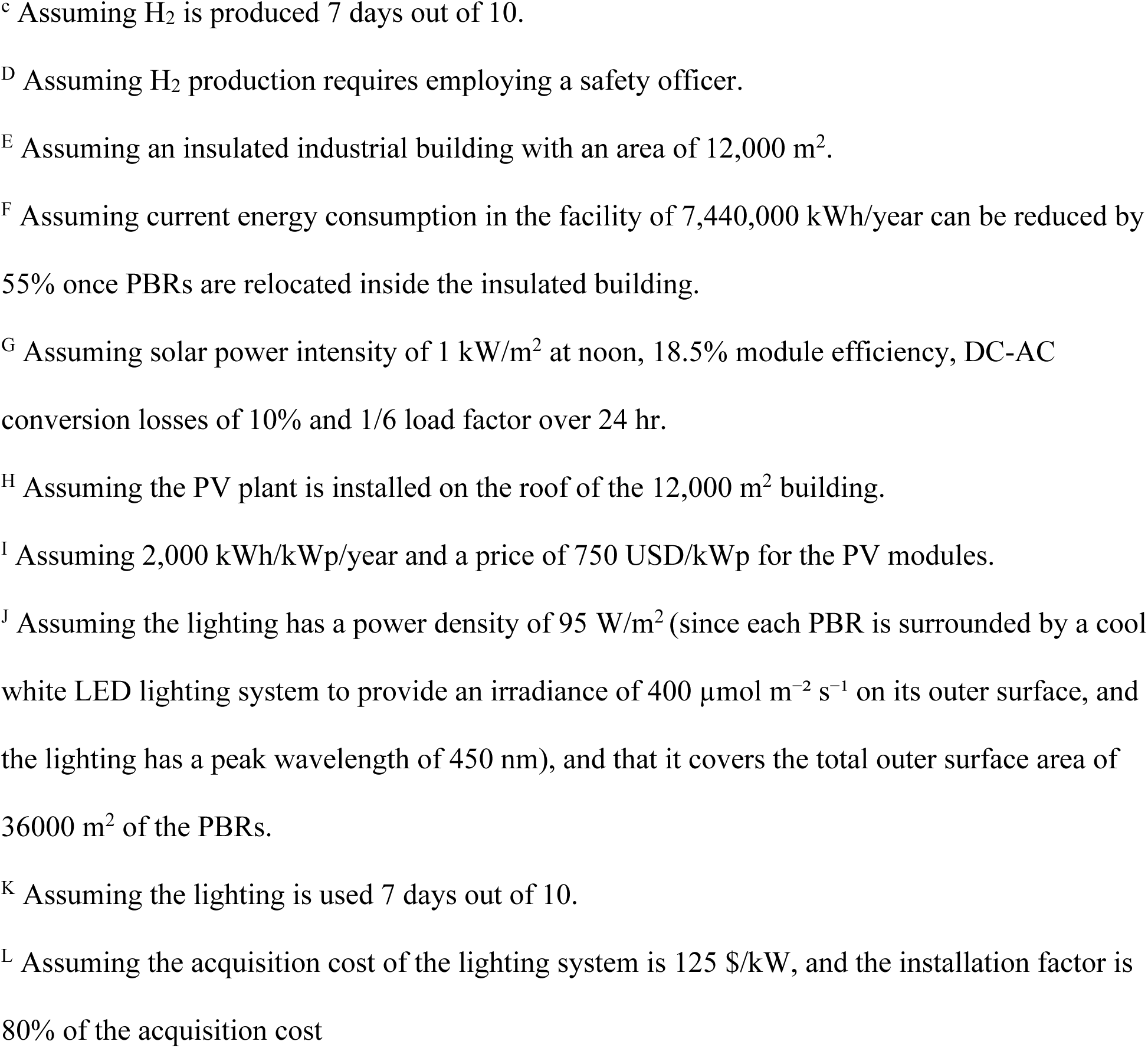

| Category | Symbol (units) | Value or equation | Ref. |
| --- | --- | --- | --- |
| Yearly H<sub>2</sub> production | $H_{2prod}$ (kg) | As detailed in the paper | |
| Onsite H<sub>2</sub> storage cost <sup>A</sup> | (\$) | $\frac{H_{2prod}}{52} \times 350$ | 43 |
| H<sub>2</sub> compression energy <sup>B</sup> | $Compr_{energy}$<br>(kWh/year) | $1.05 \times H_{2prod}$ | 44 |
| Compressor power rating <sup>c</sup> | $Compr_{power}$ (kW) | $\frac{Compr_{energy}}{2087 \times 0.7}$ | 45 |
| Compressor cost | (\$) | $47,763 \times Compr_{power}^{0.4603}$ | 45 |
| Manpower cost <sup>D</sup> | (\$/year) | 54,000 | 46 |
| Carbon credits | (\$/ton CO<sub>2eq</sub>) | 63 | 47 |
| Building cost <sup>E</sup> | (\$/m<sup>2</sup>) | 550 | 48 |
| HVAC energy savings <sup>F</sup> | (kWh/year) | $0.55 \times 7,440,000 = 4,092,000$ | 49–51 |
| PV energy output <sup>G</sup> | $PV_{out}$<br>(kWh/m<sup>2</sup>/year) | $1 \text{ kW/m}^2 \times 18.5\% \times 0.9 \times 24 \text{ h/day}$<br>$\times 365 \text{ day/yr} \times 1/6 = 240$ | |
| Total PV capacity <sup>H</sup> | $PV_{cap}$ (kWh/year) | $PV_{out} \times 12,000 \text{ m}^2$ | |
| PV cost <sup>I</sup> | (\$) | $PV_{cap} \times 2,000 \times 750$ | 52 |
| Lighting power <sup>J</sup> | $Light_{power}$ (kW) | $95 \times 36,000 = 3,447,360 \text{ W} =$<br>$3,447 \text{ kW}$ | |
| Average daily lighting duration | $Light_{duration}$<br>(h/day) | 14 (open facility) or 24 (building) | |
| Lighting energy consumption <sup>K</sup> | (kWh/year) | $Light_{power} \times Light_{duration} \times$<br>$0.7 \times 365$ | |
| Lighting system cost <sup>L</sup> | (\$) | $Light_{power} \times 125 \times 1.8 = 775,000$ | 53 |
| Energy cost | (\$/kWh) | 0.12 | 54 |
| Energy carbon footprint | (CO<sub>2eq</sub>/kWh) | 0.4 kg | 55 |

<sup>A</sup> Assuming required onsite storage volume for 1 week of production.
<sup>B</sup> Assuming H<sub>2</sub> is compressed (isothermally) from 20 bar to 350 bar prior to onsite storage.
<sup>c</sup> Assuming H<sub>2</sub> is produced 7 days out of 10.
<sup>d</sup> Assuming H<sub>2</sub> production requires employing a safety officer.
<sup>e</sup> Assuming an insulated industrial building with an area of 12,000 m<sup>2</sup>.
<sup>f</sup> Assuming current energy consumption in the facility of 7,440,000 kWh/year can be reduced by 55% once PBRs are relocated inside the insulated building.
<sup>g</sup> Assuming solar power intensity of 1 kW/m<sup>2</sup> at noon, 18.5% module efficiency, DC-AC conversion losses of 10% and 1/6 load factor over 24 hr.
<sup>h</sup> Assuming the PV plant is installed on the roof of the 12,000 m<sup>2</sup> building.
<sup>i</sup> Assuming 2,000 kWh/kWp/year and a price of 750 USD/kWp for the PV modules.

## Data availability

The authors declare that all data supporting the findings of this study are available within the paper and the supplementary file. Source data for all Figures and tables are provided with the paper.

## Acknowledgements

We Thank Dr. Doron Eisenstadt for helpful discussions and ongoing support and Eran Rosen and its team from the workshop at the Faculty of Exact Sciences, TAU. This study was financed by grants from Israel innovative authority 1737, Israel Science foundation 941/22 and VATAT Excellence center -Energy.

## Author contributions

T.E. performed all microalgal experiments and analyses. S.I. performed the technoeconomic analysis. T.E., I.Y. and S.I. wrote the manuscript. I.Y. supervised the research.

