## Supplementary material for "Photobiological Hydrogen Production at Scale: Integrating Bioprocess Optimization and Techno-Economic Modeling": (Fig. S1)

[1] School of Plant Sciences and Food Security, The George S. Wise Faculty of Life Sciences, Tel Aviv University, Aviv, Israel.

ORCIDS

Tamar Elman 0000-0003-4780-9029

Shabtai Isaac 0000-0001-8601-7601

Iftach Yacoby 0000-0003-0177-0624

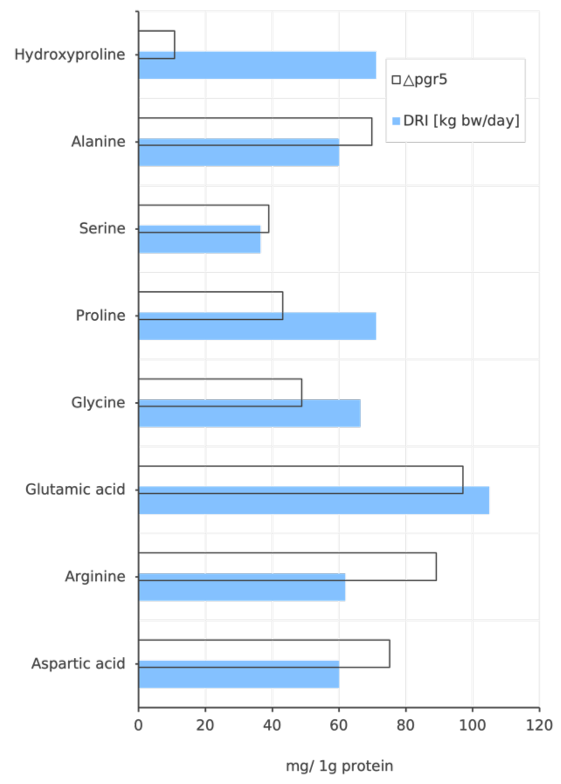

**Supplementary Figure 1. Non-Essential Amino Acid Profile of Algal Biomass After $H_2$ Production.**

Non-essential amino acid content of *pgr5* biomass after $H_2$ production is shown in milligrams per gram of protein and compared to Dietary Reference Intake values in milligrams per kilogram of body weight per day.

| Saturated fatty acids | | |
|---|---|---|
| Fatty acid | Shorthand | *pgr5* mg/ g oil |
| Lauric acid | C12 | 13.5 |
| Myristic acid | C14 | 68.8 |
| Pentadecylic acid | C15 | 11.7 |
| Palmitic acid | C16 | 3666.2 |
| Palmitic acid | C16:1 c9 | 108.9 |
| Margaric acid | C17 | 28.5 |
| Stearic acid | C18 | 1014.6 |
| Stearic acid | C18.1 trans | 81.6 |
| Stearic acid | C18.1 | 3250.1 |
| Stearic acid | C18.2 trans | 64.3 |
| Stearic acid | C18.2 cis-9,12 | 846.0 |
| Stearic acid | C18.3 trans | 276.7 |
| Stearic acid | C18.3 cis-9,12,15 ALA | 229.2 |
| Nonadecylic acid | C19.1 | 57.6 |
| Arachidic acid | C20 | 5.7 |
| Arachidic acid | C20.1 c11 | 38.4 |
| Arachidic acid | C20.3 cis-8,11,15 | 44.8 |
| Arachidic acid | C20.4 cis-5,8,11,14 | 9.1 |
| Behenic acid | C22 | 15.7 |
| Behenic acid | C22.1 c13 | 13.4 |
| Behenic acid | C22.3 cis-13,16,19 | 97.6 |
| Behenic acid | C22.4 n3 | 107.8 |
| Behenic acid | C22.5 n3 | 15.1 |
| Tricosylic acid | C23 | 4.3 |
| Lignoceric acid | C24 | 21.1 |

**Supplementary Table 1. Saturated Fatty Acids in Algal Biomass After $H_2$ Production.** Saturated fatty acids in *pgr5* biomass after $H_2$ production are presented with their common name, shorthand notation, and concentrations expressed as milligrams per gram of extracted oil.

**Elements**

| Vitamins | *pgr5* [mg/100g dw] | DRI [mg/day] Male | DRI [mg/day] Female |
|---|---|---|---|
| B1 | 0.37 | 1.2 | 1.1 |
| B2 | 2.70 | 1.3 | 1.1 |
| B3 | 5.00 | 16 | 14 |
| B5 | 0.34 | 5 | |
| B6 | 1.20 | 1.3 | |
| B12 | 0.00 | 0.0024 | |
| K | 1.80 | 0.12 | 0.09 |
| C | 23.70 | 90 | 75 |
| A | 0.01 | 0.9 | 0.7 |
| B7/B8 | 0.004 | 0.03 | |
| D3 | 0.001 | 0.015 | |
| E1 | 0.10 | 15 | |

| Micro | *pgr5* [mg/100g dw] | DRI [mg/ day] |
|---|---|---|
| Manganese | 15 | 2.3 |
| Iron | 27 | 18 |
| Copper | 20 | 0.9 |
| Zinc | 20 | 11 |
| Selenium | 0.05 | 55 |

| Macro | *pgr5* [mg/100g dw] | DRI [mg/ day] |
|---|---|---|
| Sodium | 51.0 | 1500 |
| Magnesium | 151.0 | 420 |
| Phosphorus | 807.0 | 1250 |
| Sulfur | 304.0 | 13 |
| Potassium | 246.0 | 4700 |
| Calcium | 968.0 | 1300 |

| Heavy metals | *pgr5* [mg/kg dw] | TUIL [mg/kg dw] |
|---|---|---|
| $Cd^{2+}$ | 0.08 | 5 |
| $As^{3+}$ | <0.25 | 5 |
| $Ag^{+}$ | <0.05 | NA |
| $Pb^{2+}$ | 1.7 | 5 |
| $Hg^{2+}$ | 0.2 | NA |

Color scales: [%] of DRI (0–250); [%] of TUIL (0–100)

**Supplementary Table 2. Elemental, vitamin, and heavy metal composition of pgr5 biomass following $H_2$ production.**

The table presents the concentrations of selected vitamins, minerals, and heavy metals in dried pgr5 biomass harvested after ambient $H_2$ production. Values are expressed in mg per 100 g dry weight (dw) for vitamins and minerals, and in mg/kg dw for heavy metals. Color bars indicate the percentage of the Dietary Reference Intake (DRI) met by each vitamin or mineral (capped at 250%), and the percentage of the Tolerable Upper Intake Limit (TUIL) for heavy metals (capped at 100%).

The mineral profile, shaped by the use of standardized growth media, remained consistent with previously reported values and includes particularly high levels of phosphorus (807 mg/100 g dw) and magnesium (151 mg/100 g dw), both exceeding adult daily requirements in modest portions [1]. Microalgae are widely recognized for their vitamin-rich profiles, including compounds that are rare in higher plants such as vitamins D and $B_{12}$ [2,3]. Post-$H_2$ *pgr5* biomass was found to be an excellent vitamin source. Notably, it contained exceptionally high levels of vitamin K (1.8 mg/kg dw), far exceeding levels in spinach—a commonly cited plant source (0.48 mg/kg) [4]. Significant amounts of $B_2$ (riboflavin) and $B_6$ (pyridoxine) were also detected, meeting or surpassing Dietary Reference Intakes. Notably, despite the known metal-absorbing capacity of microalgae, levels of cadmium, lead, and arsenic were well below safety thresholds [5], confirming the suitability of pgr5 biomass for nutritional and industrial applications.

| 24 hr. light | | | | Lab results | | 5-fold rate improvement | | |
|---|---|---|---|---|---|---|---|---|
| working concentration [g dw /L] | | | | Averaged rate | High rate | Averaged rate | High rate | |
| | $m^3$ | Land ($\times 1000 m^2$) | Algae (dw kg) | Kg $H_2$ | Kg $H_2$ | Kg $H_2$ | Kg $H_2$ | Kg $CO_2$ |
| 1 | 720 | 12 | 37440 | 8454 | 11913 | 42271 | 59564 | 18346 |
| 5 | 720 | 12 | 187200 | 42271 | 59564 | 211355 | 297818 | 91728 |
| 10 | 720 | 12 | 37440 | 84542 | 119127 | 422710 | 595636 | 183456 |
| Daylight (10 hr.) | | | | Lab results | | 5-fold rate improvement | | |
| working concentration [g dw /L] | | | | Averaged rate | High rate | Averaged rate | High rate | |
| | $m^3$ | Land ($\times 1000 m^2$) | Algae (dw kg) | Kg $H_2$ | Kg $H_2$ | Kg $H_2$ | Kg $H_2$ | Kg $CO_2$ |
| 1 | 720 | 12 | 37440 | 3523 | 4964 | 17613 | 24818 | 7644 |
| 5 | 720 | 12 | 187200 | 17613 | 24818 | 88065 | 124091 | 38220 |
| 10 | 720 | 12 | 37440 | 35226 | 49636 | 176129 | 248182 | 76440 |

**Supplementary Table 3. Yearly $H_2$ production and $CO_2$ fixation potential under different lighting conditions and algae concentrations.**

The calculations are based on a 12000$m^2$- facility, similar to the Bar Algae infrastructure, containing 720 $m^3$ of liquid culture. Hydrogen production is modeled using laboratory-derived rates, with an average rate of 0.35 L $H_2$/L/day (where 31 kg of algae generate 1 kg of $H_2$ per day) and a high rate of 0.5 L $H_2$/L/day (where 22 kg of algae generate 1 kg of $H_2$ per day). The projected values presented here assume a fivefold increase in production, based on previous successful upscaling from 1 L to 3 L reactors. Two lighting scenarios are considered: continuous artificial or mixed lighting, which enables hydrogen production throughout the day, and natural lighting, which is limited to an average of 10 hours per day based on sunlight availability in Israel. The assumed baseline algae concentration is 1 g dry weight (dw) per liter of culture, as observed under laboratory conditions. Hydrogen production estimates for higher concentrations (5 g/L and 10 g/L) were calculated proportionally, assuming a 1:1 scaling in production rates for both $H_2$ and $CO_2$. $CO_2$ fixation was estimated based on Figure 6, which demonstrates a fixation rate of 0.49 kg $CO_2$ per kg dw algae.

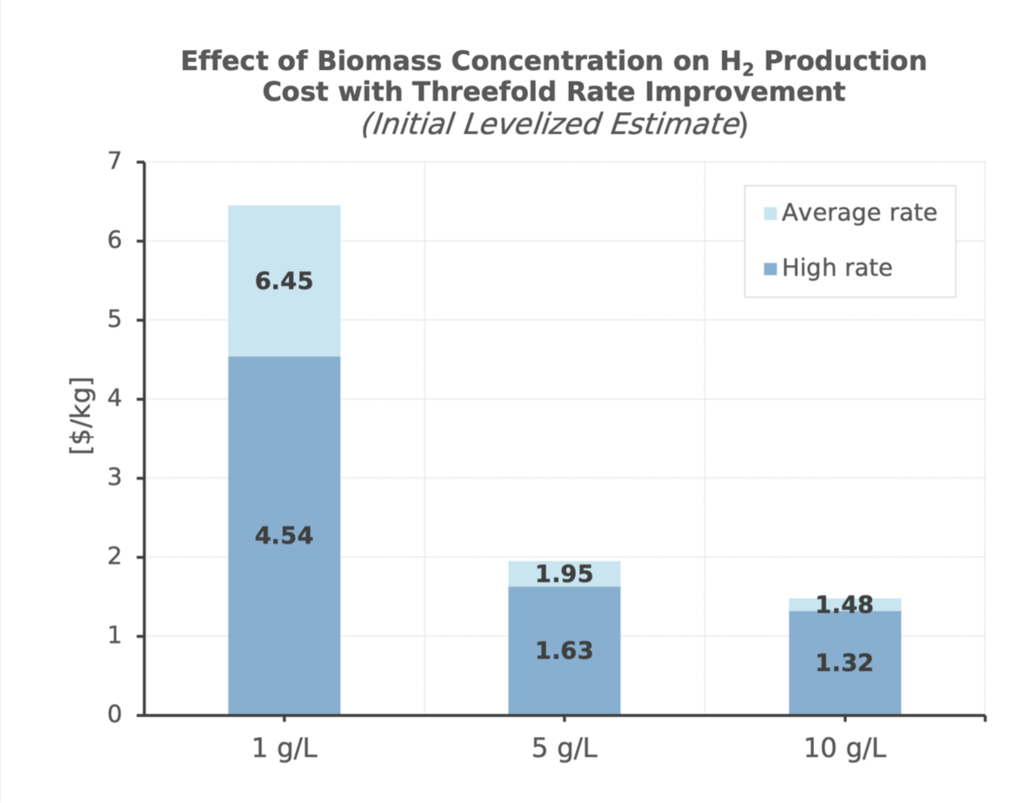

**Figure S2. Techno-economic analysis of a projected threefold improvement in hydrogen ($H_2$) production.**

Levelized $H_2$ production costs ($/kg $H_2$) under average and peak production rates at biomass concentrations of 1, 5, and 10 g/L.
